# Postsynaptic GluA3 subunits are required for the appropriate assembly of AMPA receptor GluA2 and GluA4 subunits on mammalian cochlear afferent synapses and for presynaptic ribbon modiolar-pillar morphological distinctions

**DOI:** 10.1101/2022.06.23.497300

**Authors:** Mark A. Rutherford, Atri Bhattacharyya, Maolei Xiao, Hou Ming Cai, Indra Pal, María E. Rubio

**Affiliations:** Department of Otolaryngology, Washington University School of Medicine, St. Louis, MO 63110; Department of Neurobiology, University of Pittsburgh School of Medicine, Pittsburgh, PA 15261; Department of Otolaryngology, University of Pittsburgh School of Medicine, Pittsburgh, PA 15261

**Keywords:** glutamate receptors, ultrastructure, qRT-PCR, immunolabeling, inner hear cells, 3D reconstructions, confocal imaging

## Abstract

The encoding of acoustic signals in the cochlea depends on α-amino-3-hydroxy-5-methyl-4-isoxazole propionic acid receptors (AMPARs), but relatively little is known about their reliance on specific pore-forming subunits. With 5-week-old male *GluA3*^KO^ mice, we determined cochlear function, synapse ultrastructure, and AMPAR subunit molecular anatomy at ribbon synapses between inner hair cells (IHCs) and spiral ganglion neurons (SGNs). *GluA3*^KO^ and wild-type (*GluA3*^WT^) mice reared in ambient sound pressure level (SPL) of 55-75 dB had similar ABR thresholds, wave-1 amplitudes, and latencies. Ultrastructurally, the IHC modiolar-pillar differences in presynaptic ribbon size and shape, and synaptic vesicle size seen in *GluA3*^WT^ were diminished or reversed in *GluA3*^KO^. The quantity of paired synapses (presynaptic ribbons juxtaposed with postsynaptic GluA2 and GluA4) was similar, however, GluA2-lacking synapses (ribbons paired with GluA4 but not GluA2) were observed only in *GluA3*^KO^. SGNs of *GluA3*^KO^ mice had AMPAR arrays of smaller overall volume, containing less GluA2 and greater GluA4 immunofluorescence intensity relative to *GluA3*^WT^ (3-fold difference in mean GluA4:GluA2 ratio). The expected modiolar-pillar gradient in ribbon volume was observed in IHCs of *GluA3*^WT^ but not *GluA3*^KO^. Unexpected modiolar-pillar gradients in GluA2 and GluA4 volume were present in *GluA3*^KO^. GluA3 is essential to the morphology and molecular composition of IHC-ribbon synapses. We propose the hearing loss seen in older male *GluA3*^KO^ mice results from progressive synaptopathy evident in 5-week-old mice as increased abundance of GluA2-lacking, GluA4 monomeric, Ca^2+^-permeable AMPARs.

## Introduction

In the cochlear ganglion and the ascending central auditory system, hearing relies on fast excitatory synaptic transmission via unique α-amino-3-hydroxy-5-methyl-4-isoxazolepropionic acid receptors (AMPARs) (Raman et al., 1994; Ruel et al., 1999; Gardner et al., 1999; Glowatzki and Fuchs, 2002). AMPARs are tetrameric ionotropic receptor-channels comprised of GluA1-4 pore-forming subunits plus auxiliary subunits conferring distinct electrophysiological kinetics, unique biological structures, and different pharmacological sensitivities (Jackson et al., 2011; Bowie, 2018; Azumaya et al., 2017; Twomey et al., 2018). In the adult brain, most AMPAR tetramers contain an RNA-edited form of the GluA2 subunit rendering the channel relatively impermeable to Ca^2+^, resulting in Ca^2+^-impermeable AMPARs (CI-AMPARs; Sommer et al., 1991; Higuchi et al., 1993). AMPARs lacking edited GluA2 are called Ca^2+^-permeable AMPARs (CP-AMPARs) because they have greater permeability to Ca^2+^ and larger overall ionic conductance, carried mainly by Na^+^ (Hollmann et al., 1991; Geiger et al., 1995). The expression of GluA2-lacking CP-AMPARs is downregulated in the developing brain (Pickard et al., 2000; Kumar et al., 2002; Henley and Wilkinson, 2016). However, CP-AMPARs persist or even increase with developmental maturation in some neurons of the auditory brainstem where CP-AMPARs enriched in GluA3 and GluA4 subunits are thought to be essential for fast transmission of acoustic signals (Trussell, 1997; Gardner et al., 2001; Lawrence and Trussell, 2000; Sugden et al., 2002; Wang and Manis, 2005; Youssoufian et al., 2005; Luján et al., 2019).

Cochlear afferent projections process fast auditory signals through innervation of the anteroventral cochlear nucleus, at the endbulb of Held synapses onto bushy cells, where the AMPARs are comprised mainly of GluA3 and GluA4 subunits with high Ca^2+^ permeability and rapid desensitization kinetics (Wang et al., 1998; Rubio et al., 2017). The number and distribution of fast kinetic GluA3 and GluA4 subunits in the cochlear nucleus depends on the target cells (Rubio et al., 2017). Mice lacking the GluA3 subunit have impaired auditory processing due to effects on synaptic transmission associated with altered ultrastructure of synapses between endbulbs and bushy cells (García-Hernández et al., 2017; Antunes et al., 2020). Mice lacking the GluA4 subunit have altered acoustic startle response and impaired transmission at the next synaptic relay in the midbrain at the calyx of Held, a high-fidelity central synapse (Yang et al., 2011; García-Hernández and Rubio, 2022). The rapid processing of auditory signals in the brainstem is supported by high-fidelity initial encoding of sound at synapses between cochlear inner hair cells (IHCs) and spiral ganglion neurons (SGNs) (Rutherford and Moser, 2016; Rutherford et al., 2021), however, relatively little is known about how specific pore-forming subunits affect the molecular composition of cochlear AMPARs.

In the cochlea, each primary auditory nerve fiber (i.e., SGN) is unbranched and driven to fire spikes by the release of glutamate from an individual IHC ribbon synapse acting on a single, large post-synaptic density (PSD) of approximately 850 nm in length, on average, in cat and mouse (Liberman, 1980; Payne et al., 2021). Cochlear AMPARs are comprised of GluA2-4 but not GluA1 (Niedzielski and Wenthold, 1995; Matsubara et al., 1996; Parks, 2000; Shrestha et al., 2018). Here, we examined the influence of GluA3 subunits on afferent synapse ultrastructure and on AMPAR-subunit molecular anatomy in the PSD of the auditory nerve fiber in the cochlea, with attention to GluA2 and GluA4 *flip* and *flop* isoforms and to position of innervation on the IHC modiolar-pillar axis. At the central auditory nerve projection in the cochlear nucleus, at the endbulb of Held synapse, GluA3 is required for both postsynaptic and presynaptic maturation of synapse structure and function (García-Hernández et al., 2017; Antunes et al., 2020). Therefore, we also examined presynaptic ribbon morphology in relation to position on the IHC modiolar-pillar axis, which is expected to show smaller and more spherical ribbons on the side of the IHC facing the pillar cells and the outer hair cells (pillar side) relative to the ribbons on the modiolar side facing the ganglion (Merchán-Perez and Liberman, 1996; Payne et al., 2021). Our findings in young adult male *GluA3*^KO^ mice include dysregulation of GluA2 and GluA4 subunit relative abundance and alterations in pre- and post-synaptic ultrastructure associated with an increased vulnerability to glutamatergic synaptopathy at ambient, background levels of sound.

## Materials and Methods

### Animals

A total of 26 C57BL/6 wild type (*GluA3*^WT^, n= 13) and GluA3-knockout (*GluA3*^KO^, n= 13) mice were used in this study. Generation of the GluA3^KO^ mice has been previously described (García-Hernández et al., 2017; Rubio et al., 2017). Male WT and KO mice were compared at five weeks of age (postnatal day 35 [P35]) following normal rearing in an animal facility with 55-75 dB SPL ambient noise and a 12-hour light/12-hour dark daily photoperiod. Mice were fed *ad libitum*. All experimental procedures were in accordance with the National Institute of Health guidelines and approved by the University of Pittsburgh Institutional Animal Care and Use Committee.

### Auditory Brainstem Recordings (ABR)

To test the auditory output of the *GluA3*^WT^ and *GluA3*^KO^ mice, we performed ABR as previously described (Clarkson et al., 2016; García-Hernández et al., 2017, 2022; Weisz et al., 2021). Recordings were conducted under isoflurane anesthesia in a soundproof chamber and using a Tucker-Davis Technologies (Alachua, FL) recording system. Click or tone stimuli were presented through a calibrated multi-field magnetic speaker connected to a 2-mm diameter plastic tube inserted into the ear canal. ABR were recorded by placing subdermal needle electrodes at the scalp’s vertex, at the right pinna’s ventral border, and the ventral edge of the left pinna. ABR were recorded in response to broadband noise clicks (0.1 ms) or tone pips of 4, 8, 12, 16, 24, and 32 kHz (5 ms). Stimuli were presented with alternating polarity at a rate of 21 Hz, with an inter-stimulus interval of 47.6 ms. The intensity levels used were from 80 dB to 10 dB, in decreasing steps of 5 dB. The waveforms of 512 presentations were averaged, amplified 20x, and digitalized through a low impedance preamplifier. The digitalized signals were transferred via optical port to an RZ6 processor, where the signals were band-pass filtered (0.3 – 3 kHz) and converted to analog form. The analog signals were digitized at a sample rate of ∼200 kHz and stored for offline analyses. Hearing threshold levels were determined from the averaged waveforms by identifying the lowest intensity level at which clear, reproducible peaks were visible. Wave 1 amplitudes were compared between GluA3^WT^ and GluA3^KO^ mice. For measurements of amplitudes, the peaks and troughs from the click-evoked ABR waveforms were selected manually in BioSigRZ software and exported as CSV files. The peak amplitude was calculated as the height from the maximum positive peak to the next negative trough.

### Immunohistochemistry and immunofluorescence

A total of 14 mice (GluA3^WT^ n= 7; GluA3^KO^ n=7) were anesthetized with a mixture of ketamine (60 mg/kg) and xylazine (6.5 mg/kg) and were transcardially perfused with 4% paraformaldehyde (PFA) in 0.1M phosphate buffer (PB) pH= 7.2. After 10 minutes of transcardial perfusion, cochleae and brains were removed from the skull and postfixed for 45 minutes on ice. Just after dissection, the cochleae followed perilymphatic perfusion with the same fixative through the oval window; the stapes were removed, and a hole was opened at the apex of the cochlea bone shell. After postfixation, the cochleae and brains were washed in 0.1M phosphate buffer saline (PBS).

Four cochleae (2 of each genotype) were decalcified in 10% EDTA in PBS for 24 hr, were cryoprotected in 10%, 20%, and 30% sucrose in 0.1M PBS, frozen on dry ice with tissue freezing medium (Electron Microscopy Sciences, Hatfield, PA), and stored at −20°C for up to one month. Brains were cryoprotected in the same sucrose dilutions gradient and frozen on dry ice. Cochleae were cut at a 20 um thickness section with a cryostat and were mounted on glass slides. Brains were cut with a slicing vibratome at 50-60 um thickness and collected on culture wheel plates containing 0.1M PBS. Cochlea and brain sections followed standard immunofluorescence and immunohistochemistry protocols described in Douyard et al. (2007) and Wang et al. (2011). Primary rabbit polyclonal antibodies against GluA2 (Millipore, AB1768; RRID:AB_2247874) and GluA1 (a gift from Robert J. Wenthold; Douyard et al., 2007) were used at a 1:500 dilution in 0.1M PBS. Cochlea sections followed incubation with an Alexa-594 goat-anti-rabbit secondary antibody (1:1000; Life Tech.). Brain slices followed an incubation in a biotinylated secondary antibody goat anti-rabbit (1:1000; Jackson Laboratories) in 0.1M PBS. After, brain sections were incubated in avidin-biotin-peroxidase complex (ABC Elite; Vector Laboratories; 60 min; RT), washed in 0.1M PBS, and developed with 3, 3-diaminobenzidine plus nickel (DAB; Vector Laboratories Kit; 2-5 min reaction). Sections were analyzed with an Olympus BX51 upright microscope, and digital images were captured with the CellSens software (Olympus S.L.).

The other ten cochleae (5 of each genotype) were shipped overnight to Washington University in Saint Louis in 0.1M PBS containing 5% glycerol for wholemount immunolabeling and confocal analysis of presynaptic ribbons (CtBP2/Ribeye) and postsynaptic AMPAR subunits GluA2, GluA3, and GluA4 as previously described (Jing et al., 2013; Ohn et al., 2016; Sebe et al., 2017; Kim et al., 2019; Hu et al., 2020). Primary antibodies: CtBP2 mouse IgG1 (BD Biosciences 612044; RRID:AB_399431), GluA2 mouse IgG2a (Millipore MAB397; RRID:AB_2113875), GluA3 goat (Santa Cruz Biotechnology SC7612), and GluA4 rabbit (Millipore AB1508; RRID:AB_90711) were used with species-appropriate secondary antibodies conjugated to Alexa Fluor (Life Tech.) fluorophores excited by 488, 555, or 647 nm light in triple-labeled samples mounted in Mowiol. Samples were batch processed using the same reagent solutions in two cohorts, including WT and KO mice.

### Confocal Microscopy and Image Analysis

For synapse counts and measurements of intensity, volume, sphericity, and position confocal stacks were acquired with a Z-step of 0.37 µm and pixel size of 50 nm in X and Y on a Zeiss LSM 700 with a 63X 1.4 NA oil objective lens. To avoid saturation of pixel intensity and to enable comparisons across images and genotypes, we first surveyed the samples to determine the necessary laser power and gain settings to collect all of the images. Then, using identical acquisition settings for each sample, we collected 3–4 images from each cochlea at each of the 3 cochlear regions (basal, middle and apical, respectively) centered near tonotopic characteristic frequencies of 10, 20, and 40 kHz (Müller et al., 2005). The numbers of hair cells and paired and unpaired pre- and post-synaptic puncta were counted and manually verified after automated identification using Imaris software (Bitplane) to calculate the mean per IHC per image. The experimenter was blinded to the mouse genotype. For each group of images from which synapses were counted or synaptic properties were measured, grand means (±SD) were calculated across image means (Fig. 6B; 7E,G; 8A-D). Paired synapses were identified as juxtaposed puncta of presynaptic ribbons (CtBP2) and postsynaptic AMPARs (GluA2 and/or GluA4), which appear to partly overlap at confocal resolution (Rutherford, 2015). Unpaired (i.e., lone) ribbons were defined as CtBP2 puncta in the IHC but lacking appositional GluA2 or GluA4 puncta. For unpaired ribbons, we did not distinguish membrane-anchored from unanchored. Ribbonless synapses consisted of GluA2 and/or GluA4 puncta located around IHC basolateral membranes but lacking CtBP2. Pixels comprising puncta of synaptic fluorescence were segmented in 3D as ‘surface’ objects in Imaris using identical settings for each image stack, including the ‘local contrast background subtraction’ algorithm for automatically calculating the threshold pixel intensity for each fluorescence channel in each image. This adaptive and automatically-calculated thresholding algorithm compensated for differences in overall luminance between image stacks that would affect the volume of segmented puncta if a fixed threshold was applied across images, and avoided the potential subjective bias of setting a user-defined arbitrary threshold value separately for each image. Intensity per synaptic punctum was calculated as the summation of pixel intensities within the surface object. To associate the intensities of different GluA puncta belonging to the same synapse (Fig. 7C; Fig. 9E-H), we generated surface objects from a virtual 4^th^ channel equal to the sum of the three channels (GluA2, 3, and 4; or CtBP2, GluA2, and GluA4) and then summated the pixel intensities within each of the three fluorescence channels comprising each synapse defined as a punctum on the 4^th^ channel. The mean density of synaptic fluorescence per image (Fig. 8C) was calculated as mean punctum Intensity (a.u.) divided by mean punctum Volume (µm^3^) using surface objects calculated from corresponding individual fluorescence channels. To associate the volumes of different GluA puncta belonging to the same synapse (Fig. 7B), we used the virtual 4^th^ channel to generate masks. The mask for each synapse had a unique color value. Objects belonging to the same synapse were identified based on common overlap with the unique color value assigned to each mask. Sphericity is the ratio of the surface area of a sphere to the surface area of an object of equal volume, so sphericity of 1 means the object is a perfect sphere. To differentiate synapse position, images were used in which the row of IHCs was oriented with the modiolar-pillar dimension in the microscope’s Z-axis and the organ of Corti was not sloping in the image volume, and modiolar-side and pillar-side groups were split at the midpoint of the range of synapses along the modiolar-pillar dimension.

### Reverse transcription-polymerase chain reaction (RT-PCR) and quantitative PCR (qPCR)

Under isofluorane anesthesia, mice (*GluA3*^WT^ n= 4; *GluA3*^KO^ n= 4) were euthanized via cervical dislocation and decapitation. Immediately after the following decapitation, the cranium was opened, and the inner ears were removed. Inner ears were flash-frozen in liquid nitrogen and stored in −80°C for up to one month until RT-PCR. In preparation for RT-PCR, both inner ears from each individual were homogenized by hand with mortar or pestle and RNA was extracted with Trizol (Ambion by life technology). The RNA pellet was resuspended and the supernatant containing RNA from each individual’s inner ears was prepared for RT-PCR using the SuperScript Strand Synthesis System kit (Invitrogen, cat. no. 11904018). The resulting cDNA was stored at −20°C for one week or less before real-time qPCR. qPCR was performed at the Genomics Research Core at the University of Pittsburgh using EvaGreen qPCR kit (MidSci, Valley Park, MO, cat. no. BEQPCR_R) and primers for *Gria2* and *Gria4 flip* and *flop*, which were the same primers used successfully in a previous RT-qPCR experiment by Hagino et al. (2004). In a 25 µl PCR reaction mixture, 2 µl cDNA samples were amplified in a Chromo 4 detector (MJ Research, Waltham, MA). GAPDH and 18S rRNA were used as housekeeping genes. Each sample (consisting of RNA product of both cochleae from each mouse) was run in triplicate, and average cycle thresholds (CTs) were used for quantification. Relative abundances of each splice isoform for *GluA3*^KO^ males compared to *GluA3*^WT^ were reported as fold-change, calculated using the following equation: 2Delta-CT (^2-ΔΔ^CT), where ^ΔΔ^CT = (CT_GluA3_^WT^– CT_GAPDH_ or CT_18S rRNA_) – (CT ^KO^ – CT or CT), and CT represents the cycle threshold of each cDNA sample. For a more in-depth explanation of this equation see Schmittgen and Livak (2008). Electrophoresis of 10 µl of RT-PCR products was performed using 3% agarose (SeaKem LE Agarose by Lonza) with molecular ladder gel containing 0.5 µg/ml ethidium bromide in ^x^0.5 tris-acetate-ethylenediaminetetraacetic acid (TAE) buffer (pH: 8.0) and run at 100 V for 60 min. The size and thickness of the agarose gel, reagents, and other conditions were kept constant. The band-size and DNA concentration of each PCR amplicon was determined by comparison to the corresponding band in the molecular weight ladder (Gene Ruler 100 BP DNA ladder Thermo Scientific). The amplicon images (RT-PCR bands) in the gel were captured under ultraviolet (UV) light and documented using a Bio Rad Molecular Imager Gel Doc RX+ Imaging system. All the parameters and experimental conditions used were kept constant throughout the study. The image was saved (in JPEG format) on a computer for digital image analysis using ImageJ software. The mean gray value (MGV) of each band was determined with NIH-ImageJ software (https://imagej.nih.gov/ij/). Samples were normalized to GAPDH. The *flip/flop* ratio was obtained by dividing the MGVs of the *flip* by the *flop*.

### Transmission Electron Microscopy (TEM)

Four mice (2 per genotype) were anesthetized with a mixture of ketamine (60 mg/kg) and xylazine (6.5 mg/kg) and were transcardially perfused with 0.1M PB, followed by 3% PFA and 1.5% glutaraldehyde in 0.1 M PB. Cochleae were dissected from the temporal bones, and fixative was slowly introduced through the oval window after removing the stapes and opening a hole at the apex of the cochlea bone shell. Cochleae were post-fixed overnight in the same fixative at 4°C and followed a protocol slightly modified from Clarkson et al. (2016). After decalcification in 10% EDTA for 24 hours at 4°C on a rotor, cochleae were washed in 0.1M cacodylate buffer and postfixed with 1% osmium and 1.5% potassium ferrocyanide in cacodylate buffer for 1 hr at room temperature (RT). Cochleae were dehydrated in an ascending ethanol gradient (ETOH; 35%, 50% 70%, 80% 90%) and were blocked-stained with 3% uranyl acetate in 70% ETOH for 2 hr at 4°C before the 80% ETOH. The latest dehydration steps performed with 100% ETOH and propylene oxide were followed by infiltration with epoxy resin (EMBed-812; Electron Microscopy Science, PA USA). Cochleae were cut with a Leica EM UC7 ultramicrotome, and series of 15-20 serial ultrathin sections (75-80 nm in thickness) were collected. Each serial ultrathin section was collected on numbered single slot gold-gilded grids with formvar. Ultrathin sections were observed with a JEOL-1400 transmission electron microscope (TEM; JEOL Ltd., Akishima Tokyo, Japan), and images of the midcochlea (∼20 kHz) containing inner hair cell (IHC)-ribbon synapses of the modiolar and pillar side were captured with an Orius^TM^ SC200 CCD camera (Gatan Inc., Warrendale, PA, USA).

### Three-Dimensional (3-D) Reconstructions and NIH Image-J Analysis of TEM micrographs

TEM micrographs (at x40,000 magnification) of the serial IHC-ribbon synapses were aligned and reconstructed using the Reconstruct software (https://synapseweb.clm.utexas.edu/software-0; Fiala, 2005) as previously described (Gómez-Nieto and Rubio, 2009; Clarkson et al., 2016, 2020). A total of 27 (*GluA3*^WT^) and 30 (*GluA3*^KO^) IHC-ribbon synapses were reconstructed. In brief, two successive sections were aligned via rotation and translation such that corresponding structures like mitochondria in the two sections were superimposed. Linear transformation compensated for distortions introduced by the sectioning. Following alignment of the TEM sections, structures of interest were segmented visually into contours of separate objects. The subsequent linear interpolation between membrane contours in adjacent images resulted in polygonal outlines of cell membranes, synaptic vesicles, and synaptic ribbons. The 3-D rendering was generated as VRML files from the stacks of all contoured sections. We calculated volumes and surface areas of the structures of interest by filling these stacks with tetrahedra. A total of 44 (*GluA3*^WT^) and 52 (*GluA3*^KO^) single TEM micrographs (at x40,000 magnification) were used for analysis of the synaptic ribbon major axis and circularity as well as synaptic vesicles (SV) size using NIH Image-J software.

### Statistical Analysis

Statistical analysis was performed with GraphPad Prism software (Version 9.3.1) or with IGOR Pro software (Wavemetrics, Version 7.08). Mann Whitney or simple t-tests were used to compare two independent groups. One-way ANOVA or Kruskal-Wallis tests and two-way ANOVA were used for comparisons in which there were one and two independent variables, respectively. Paired and multiple comparisons were made using Šidák’s and Tukey’s tests, respectively. Statistical significance for all tests was set to *p* < α; α = 0.05. Data are represented as mean ± standard deviation (SD). The coefficient of variation (CV) was calculated as CV = SD / mean.

## Results

### Hearing sensitivity is unaltered in 5-week-old *GluA3*^KO^ mice; transcription and mRNA splicing of GluA2 and GluA4 isoforms are similar in cochleae of *GluA3*^WT^ and *GluA3*^KO^

We first determined whether the cohort of 5-week-old C57BL/6J *GluA3*^WT^ and *GluA3*^KO^ differed in their auditory sensitivity. Our ABR analysis showed no differences between genotypes in clicks or pure tone thresholds or wave-1 amplitude or latency (Fig. 1A). We note that male *GluA3*^KO^ and *GluA3*^WT^ mice at two months of age have similar ABR thresholds but *GluA3*^KO^ mice have reduced ABR wave-1 amplitude (García-Hernández et al., 2017), suggesting cochlear deafferentation sometime between ages P35 and P60. We then asked if GluA3 was required for the appropriate expression of GluA1-4 subunits in the cochlear spiral ganglion (the auditory nerve fiber somata) and in the cochlear nucleus (Fig. 1B-C). In WT mice, mature SGNs express GluA2, GluA3, and GluA4 subunits of the AMPAR, but not GluA1 (Niedzielski and Wenthold, 1995; Matsubara et al., 1996; Parks, 2000; Shrestha et al., 2018). With immunolabelling, we observed GluA2 in the SGNs of both genotypes (Fig. 1B**, upper left**). We found that SGNs lacked GluA1 in *GluA3*^WT^ mice, as expected, and we did not observe compensatory GluA1 expression in SGNs of *GluA3*^KO^ (Fig. 1B**, lower left**). We also checked the immunolabeling of GluA1 on brainstem sections containing the ventral cochlear nucleus and cerebellum (Fig.1B**, right**). As expected, we found GluA1 immunoreactivity in the cerebellar Bergmann glia of *GluA3*^WT^ and *GluA3*^KO^ mice (Matsui et al., 2005; Douyard et al., 2007). In contrast, as in the SGNs, the ventral cochlear nucleus of 5-week-old mice lacked GluA1 immunoreactivity in *GluA3*^WT^, as previously shown (Wang et al., 1998), and we discovered no compensation in *GluA3*^KO^. At ribbon synapses in the cochlea, the PSDs on the postsynaptic terminals of SGNs expressed GluA2, 3, and 4 in *GluA3*^WT^ as previously shown (Sebe et al., 2017), while those in *GluA3*^KO^ lacked immunolabeling for GluA3 (Fig. 2). This confirmed the deletion of GluA3 subunits was effective in SGNs of *GluA3*^KO^ mice, as previously shown in the cochlear nucleus (García-Hernández et al., 2017; Rubio et al., 2017), and was not associated with compensatory upregulation of GluA1 subunits.

**Figure 1.**
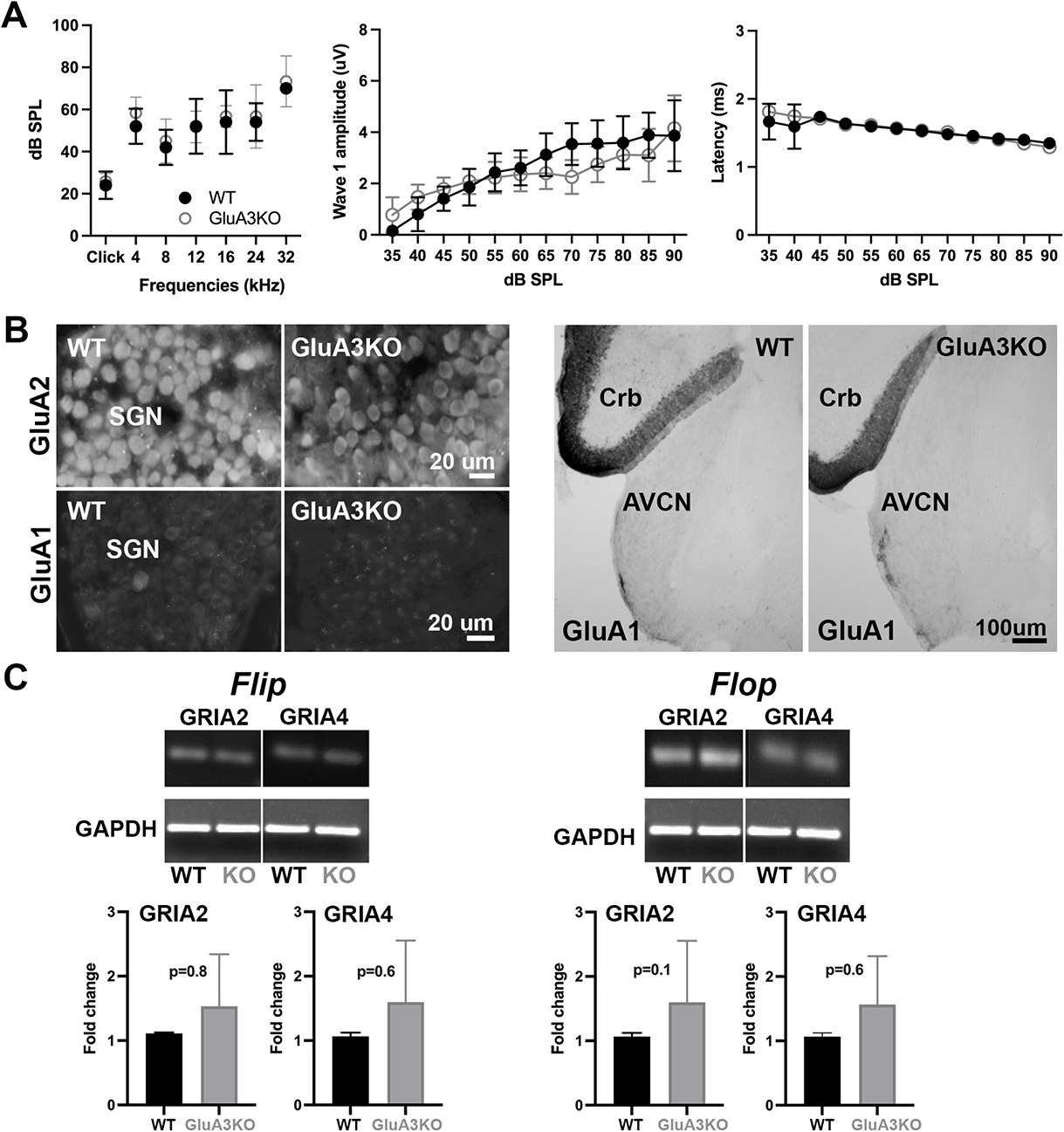
ABRs, GluA1 and GluA2 immunolabeling and qRT-PCR in *GluA3*^WT^ and *GluA3*^KO^. **A.** Mean ABR thresholds (±SD) are similar between male *GluA3*^WT^ and *GluA3*^KO^ mice (F_(1, 63-)_ = 1.599, *p*= 0.2107; RM two way-ANOVA; *GluA3*^WT^ n= 8; *GluA3*^KO^ n= 10). In WT and *GluA3*^KO^ there was a main effect of frequency (F_(6, 63)_ = 23.92, *p* < 0.0001). Mean wave 1 amplitudes were similar between *GluA3*^WT^ and *GluA3*^KO^ mice (F_(1, 132)_ = 2.419, *p*= 0.1223; RM two way-ANOVA). In both genotypes, there was an effect of sound intensity (F_(11, 132)_ = 22.65; *p*< 0.0001). Paired multiple comparisons show that only at 70 dB SPL, wave 1 amplitude is smaller in the *GluA3*^KO^ (*p*= 0.045). Mean wave latencies (±SD) are similar between *GluA3*^WT^ and GluA3^KO^ mice (F_(1, 132)_ = 0.8907, *p*= 0.3470; RM two way-ANOVA). There was an effect of sound level on wave 1 latencies (F_(1, 132)_ = 24.31, *p*< 0.0001), with significantly longer latencies at each decreasing sound level. **B.** Micrographs show immunolabeling for GluA2 and GluA1 on SGN, and for GluA1 on the anteroventral cochlear nucleus (AVCN) and cerebellum (Crb) of *GluA3*^WT^ and *GluA3*^KO^ mice. Immunolabeling for GluA2 is observed on SGN of WT and KO mice. In contrast, immunolabeling for GluA1 is not observed on SGN and AVCN of both *GluA3*^WT^ and *GluA3*^KO^ mice, but is observed in the cerebellum labeling the cerebellar Bergmann glia of both genotypes. **C.** Images of *Gria2* and *Gria4 flip* and *flop*, and GAPDH gels of *GluA3*^WT^ and *GluA3*^KO^ inner ears. Histograms show fold change (±SD) of qRT-PCR product. Independent samples *t*-test.

**Figure 2.**
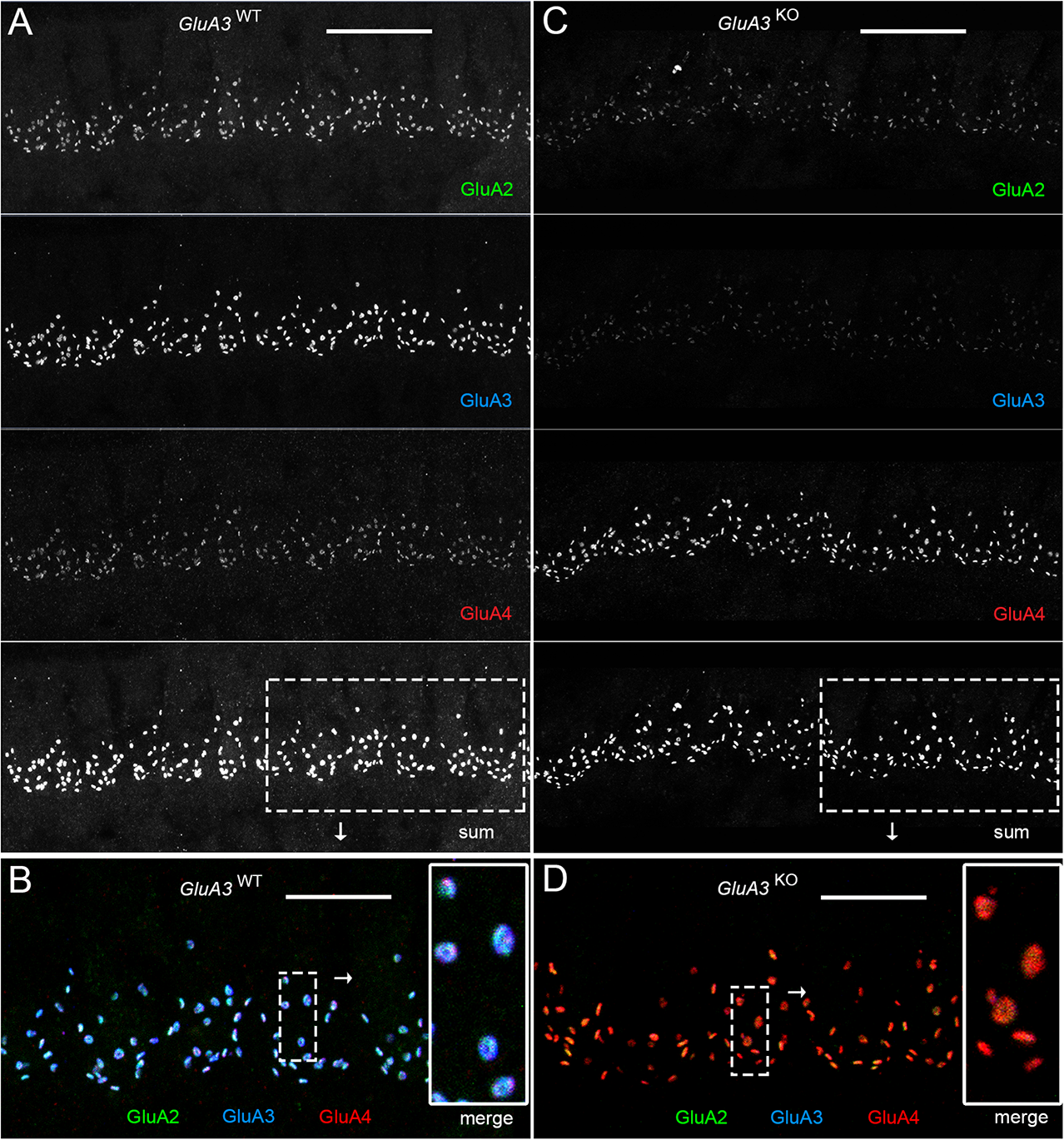
Immunohistofluorescence of AMPAR pore-forming subunits GluA2, 3, and 4 on spiral ganglion neuron postsynaptic terminals in the organ of Corti. Confocal microscope immuno-fluorescence images of afferent ribbon synapses in organ of Corti whole-mount samples from *GluA3*^WT^ (left) and *GluA3*^KO^ mice (right) in the mid-cochlea. Anti-GluA2 (green), - GluA3 (blue), and -GluA4 (red) label the postsynaptic AMPAR subunits encoded by the *Gria2, Gria3, and Gria4* genes, respectively. Each subpanel displays synaptic puncta of approximately 12 inner hair cells. Scale bars: 20 µm (A, C); 10 µm (B, D). **A.** From top to bottom: *GluA3*^WT^ in grayscale for GluA2, 3, 4, and the sum of the three. **B.** Merged color image of the region of interest indicated by the dashed rectangle in panel A. Inset on right: enlargement of the dashed rectangular region of interest on left shows 5 postsynaptic AMPA-receptor arrays of ribbon synapses from one IHC. **C.** From top to bottom: *GluA3*^KO^ in grayscale for GluA2, 3, 4, and the sum of the three. **D.** Merged color image of the region of interest indicated in panel C. Inset: enlargement of a rectangular region of interest shows several postsynaptic AMPA-receptor arrays of ribbon synapses from one IHC.

Unique isoforms are generated by alternative splicing of the pore-forming GluA subunits. In the brain, *flip* and *flop* splice variants are expressed in distinct but partly overlapping patterns and impart different desensitization kinetics (Sommer et al., 1990). The chicken and rat cochlear nuclei express predominantly the fast-desensitizing *flop* isoforms (Schmid et al., 2001; Sugden et al., 2002). With qRT-PCR, we determined whether the absence of GluA3 altered posttranscriptional *flip* and *flop* splicing of mRNA for GluA2 (*Gria2*) or GluA4 (*Gria4*) in the cochlea. Comparing the levels of *flip* or *flop* for *Gria2* or *Gria4* between *GluA3*^WT^ and *GluA3*^KO^, we found no significant differences (Fig. 1C). In addition, using the PCR gels, we calculated the *flip*/*flop* ratios for *Gria2* and *Gria4* in *GluA3*^WT^ and *GluA3*^KO^ and found no differences between genotype (*Gria2 flip*/*flop* ratio WT: 0.7, *GluA3*^KO^: 0.6; *Gria4 flip*/*flop* ratio, WT: 0.7, *GluA3*^KO^: 0.6).

Overall, this shows that lack of GluA3 did not affect hearing sensitivity at 5-weeks of age, in contrast to 8 weeks when ABR peak amplitudes are reduced (García-Hernández et al., 2017). Taken together with previous work, this suggests the 5-week-old *GluA3*^KO^ cochlea may be in a pathological but pre-symptomatic, vulnerable state. The levels of *Gria2* or *Gria4 flip* or *flop* mRNA isoforms in cochleae of male mice at 5-weeks of age were similar in *GluA3*^WT^ and *GluA3*^KO^, and in both genotypes, the expression of the *flop* splice variant for *Gria2* and *Gria4* is predominant.

### Pre- and post-synaptic ultrastructural features of Inner Hair Cell ribbon synapses are disrupted in the organ of Corti of *GluA3*^KO^ mice

Based on the similarity of auditory sensitivity in male *GluA3*^WT^ and *GluA3*^KO^ mice at 5-weeks of age, we hypothesized that the ultrastructure of IHC-ribbon synapses would be unaltered in the *GluA3*^KO^ mice. Qualitatively, the sensory epithelium’s general structure and cellular components were like WT and similar to published data of C57BL/6 mice (not shown; Ohlemiller and Gagnon, 2004). Synapses from the midcochlea of both *GluA3*^WT^ and *GluA3*^KO^ mice had electron-dense pre- and post-synaptic membrane specializations and membrane-associated presynaptic ribbons (Figs. 3 and 4).

**Figure 3.**
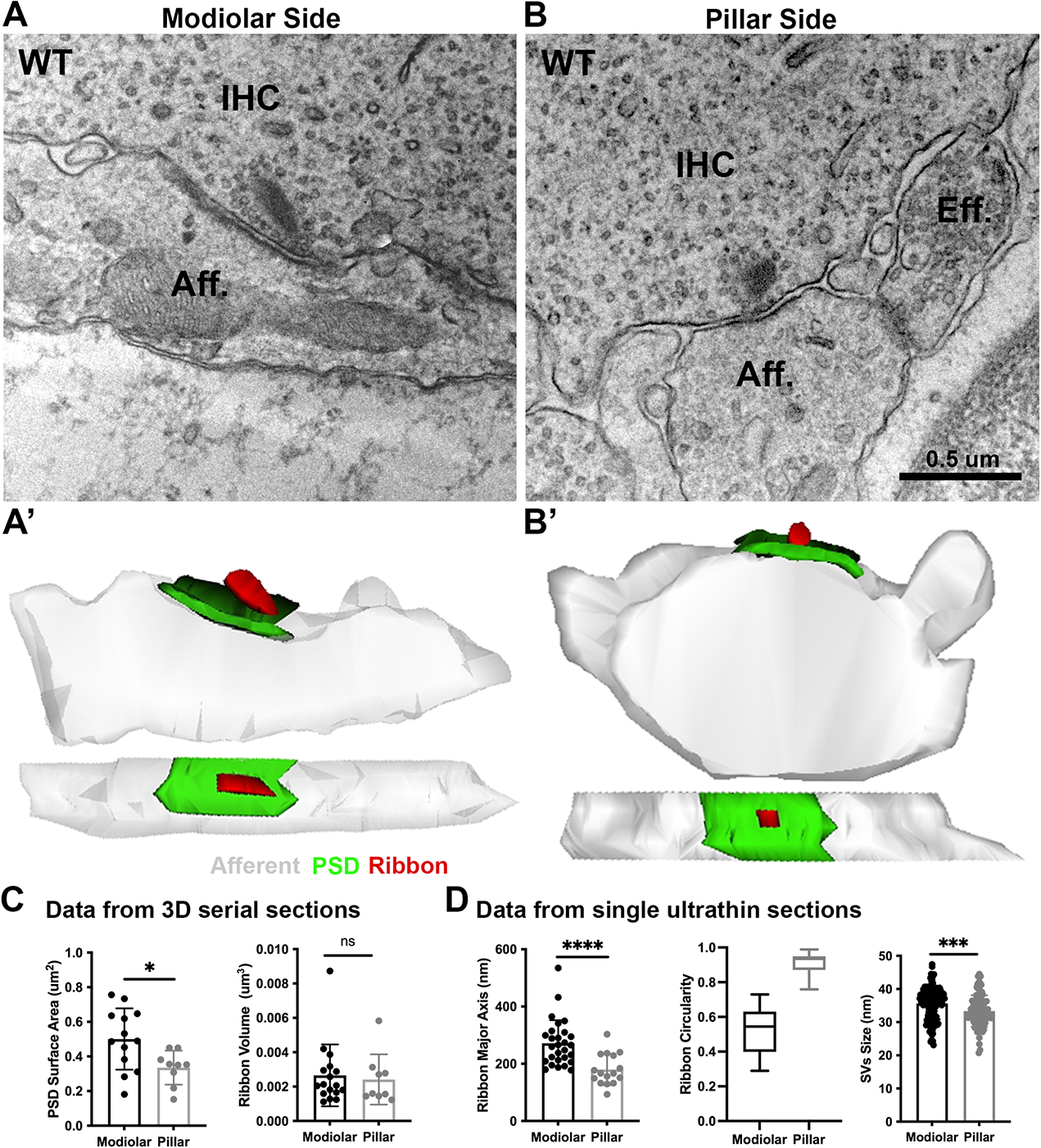
Ultrastructural features of *GluA3*^WT^ IHC-ribbon midcochlear synapses. **A-B**. TEM micrographs of IHC-synapses on the modiolar (A) and pillar side (B). Aff.: afferent; IHC: inner hair cell; Eff.: efferent terminal. **A’-B’**. Three-D reconstructions of the IHC-ribbon synapses are shown in A and B. **C**. Plots of the quantitative data of the PSD surface area and ribbon volume obtained from the 3D reconstructions of *GluA3*^WT^ mice. **D**. Plots of the quantitative data from single ultrathin sections of the major axis and circularity of the ribbons and the size of SVs of *GluA3*^WT^ mice.

**Figure 4.**
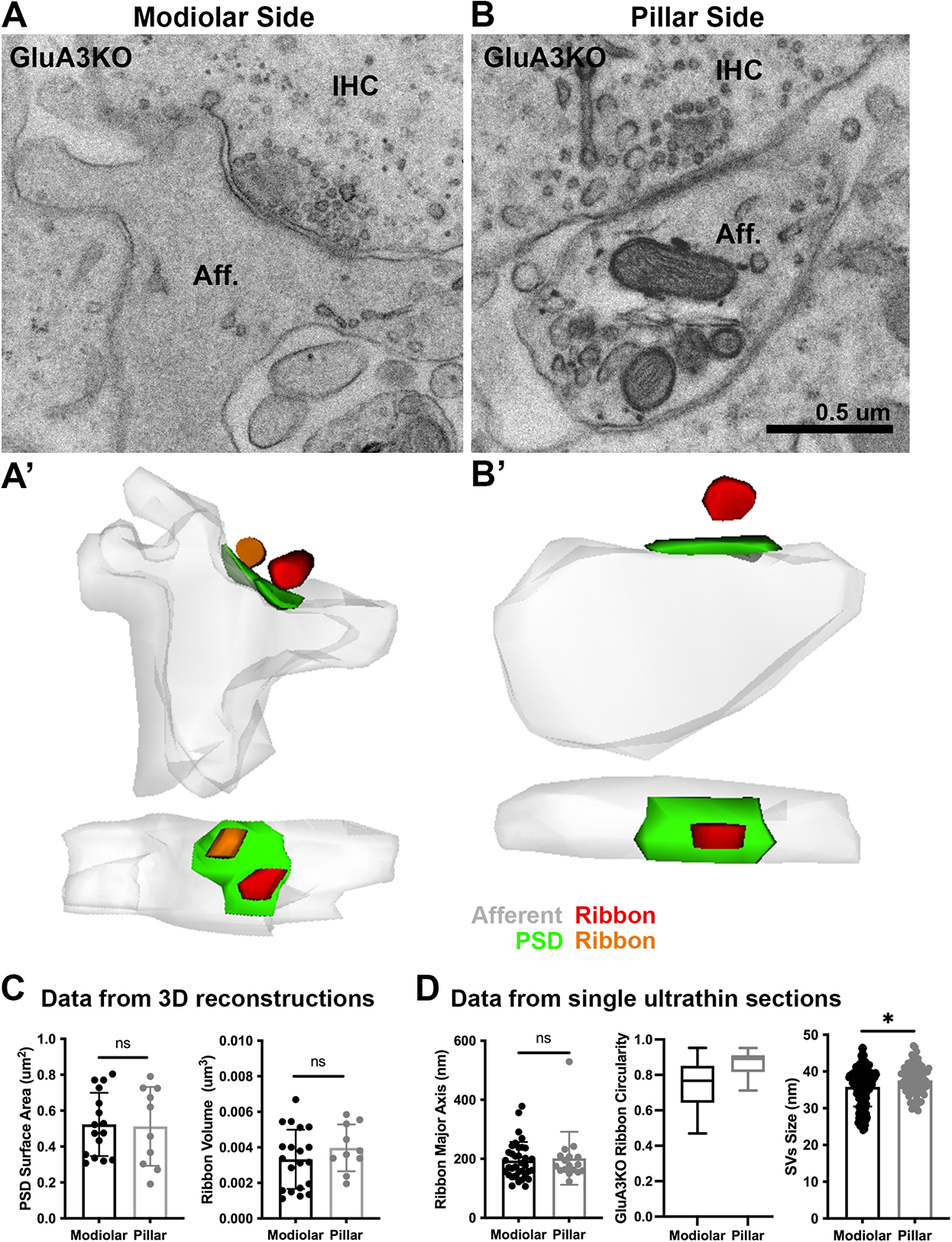
Ultrastructural features of *GluA3*^KO^ IHC-ribbon midcochlear synapses. **A-B**. TEM micrographs of IHC-synapses on the modiolar (A) and pillar side (B) of *GluA3*^KO^ mice. Aff.: afferent; IHC: inner hair cell; ER.: endoplasmic reticulum/swelling. **A’-B’**. Three-D reconstructions of the IHC-ribbon synapses shown in A and B. **C**. Plots of the quantitative data of the PSD surface area and ribbon volume obtained from the 3D reconstructions of GluA3^KO^ mice. **D**. Plots of the quantitative data from single ultrathin sections of the major axis and circularity of the ribbons and the size of SVs of *GluA3*^KO^ mice.

A total of 27 synapses of *GluA3*^WT^ mice were analyzed in 3D using serial sections (on average, 7 ultrathin sections per PSD). Of this total, 18 were on the modiolar side and 9 on the pillar side of the IHCs (Fig. 3A-B). We found the PSD surface area to be larger (*p*= 0.017, Mann-Whitney U test, U: 23) for synapses on modiolar side (mean: 0.50 ± 0.1 µm^3^) when compared to the pillar side (mean: 0.33 ± 0.09 µm^3^) (Fig. 3C**, left**). Presynaptic ribbon volume was similar between these two groups (*p*= 0.525, Mann-Whitney U test, U: 64; modiolar mean: 0.003 ± 0.001 µm^3^; pillar mean: 0.002 ± 0.001 µm^3^; Fig. 3C**, right**). Six of 18 modiolar-side synapses had 2 ribbons (33.3%), whereas all the synapses on the pillar side had only 1 ribbon. Next, a total of 44 single TEM micrographs were analyzed to measure the major axis and circularity of the ribbon (n= 46 [2 PSDs had 2 ribbons]) and the size of the SVs (n= 249). Analysis of *GluA3*^WT^ showed that the IHC-synapses on the modiolar side had longer major ribbon axes (mean: 272 ± 81 nm, CV: 0.30) and less circularity (mean: 0.5 ± 0.1) compared to the pillar-side ribbons (mean major axes: 180 ± 55 nm, CV: 0.30; mean circularity: 0.9 ± 0.01). These data show that ribbons on the modiolar side of *GluA3*^WT^ IHCs are elongated, while those on the pillar side are more round in shape (Fig. 3D**, left and center**), as previously shown for C57BL/6 mice at 5-weeks of age (Payne et al., 2021). Analysis of SV size showed that the SVs of modiolar-side synapses were larger (*p*= 0.0001, Mann-Whitney U test, U: 4975) than those of the pillar-side synapses (modiolar: 36 ± 5 nm, CV: 0.13; pillar: 33 ± 4 nm, CV: 0.14; Fig. 3D**, right**).

From *GluA3*^KO^, a total of 30 synapses were analyzed in 3D with serial sections (on average, 7 ultrathin sections per PSD). Of this total, 20 were on the modiolar side and 10 on the pillar side of the IHCs (Fig. 4A-B). Analysis showed that PSD surface area and ribbon volume were similar for modiolar- and pillar-side synapses (PSD surface area *p*= 0.72, Mann-Whitney U test, U: 73; ribbon volume *p*= 0.52 Mann-Whitney U test, U: 64) (mean PSD surface area, modiolar: 0.52 ± 0.17 µm^3^, pillar: 0.51 ± 0.22 µm^3^; mean ribbon volume, modiolar: 0.003 ± 0.001 µm^3^, pillar: 0.003 ± 0.001 µm^3^; Fig. 4C). Six of the 20 modiolar-side synapses had 2 ribbons (30%), whereas all synapses on the pillar side had only 1 ribbon. A total of 52 single TEM micrographs were analyzed to measure the major axis and circularity of the ribbon (n=52) and the size of the SVs (n=279). Analysis of *GluA3*^KO^ showed that IHC-synapses on the modiolar side had similar major ribbon axes (194 ± 63 nm, CV: 0.33) and similar circularity (0.75 ± 0.14) compared to the pillar-side ribbons (major axis: 202 ± 89 nm, CV: 0.44), circularity: 0.90 ± 0.07). Thus, unlike *GluA3*^WT^, modiolar- and pillar-side synapses had ribbons of similar size and roundedness in *GluA3*^KO^ (Fig. 4D**, left and center**). Also, opposite to the pattern in *GluA3*^WT^, SVs of modiolar-side synapses were smaller (*p* = 0.022, Mann-Whitney U test, U: 7030) than those of pillar-side synapses in *GluA3*^KO^ (modiolar: 36 ± 5 nm, CV: 0.15; pillar: 38 ± 4 nm, CV: 0.10; Fig. 4D**, right**).

### Inner Hair Cell modiolar-pillar differences are eliminated or reversed in *GluA3*^KO^

We then compared PSDs and ribbons among *GluA3*^WT^ and *GluA3*^KO^ mice on the modiolar and pillar sides (Fig. 5A). Overall PSD surface area was significantly different between genotypes (*p* = 0.027, Kruskal-Walli’s test). Considering modiolar-side and pillar-side synapses separately with multiple (paired) comparisons revealed the PSD surface areas of modiolar-side synapses to be similar among *GluA3*^KO^ and *GluA3*^WT^ (*p* = 0.91). In contrast, the mean PSD surface area of the pillar-side synapses was larger in *GluA3*^KO^ than *GluA3*^WT^ (*p* = 0.007) (Fig. 5A**, left**). Synaptic ribbon volume differed between *GluA3*^WT^ and *GluA3*^KO^ (*p*< 0.0001, one-way ANOVA). Comparison analyses showed that the ribbon volumes of modiolar-side synapses were similar between *GluA3*^WT^ and *GluA3*^KO^ (*p*= 0.99). In contrast, the pillar-side synapses were larger in *GluA3*^KO^ than *GluA3*^WT^ (*p* = 0.0006) (Fig. 5A**, right**). Differences between the ribbon major axis were found between WT and KO (*p*< 0.0001; one-way ANOVA). On the modiolar side, analysis of the ribbon major axis length showed that those of the *GluA3*^KO^ were significantly smaller than *GluA3*^WT^ (*p* < 0.0001), whereas pillar-side synapses were similar in major axis length (Fig. 5B**, left**) (*p* > 0.5). Differences in ribbon circularity were also found between genotypes (*p* < 0.0001, one-way ANOVA). Paired comparisons showed that modiolar-side ribbons were significantly less circular in *GluA3*^WT^ (*p* < 0.0001), whereas pillar-side ribbons were of similar circularity (*p* = 0.31) (Fig. 5B**, center**). SV size differed between genotypes (*p* < 0.0001, one-way ANOVA). Data showed that SVs of modiolar synapses were similar among genotypes (*p* = 0.94), while those of pillar synapses were significantly larger in *GluA3*^KO^ (*p* < 0.0001) (Fig. 5D**, right**).

**Figure 5.**
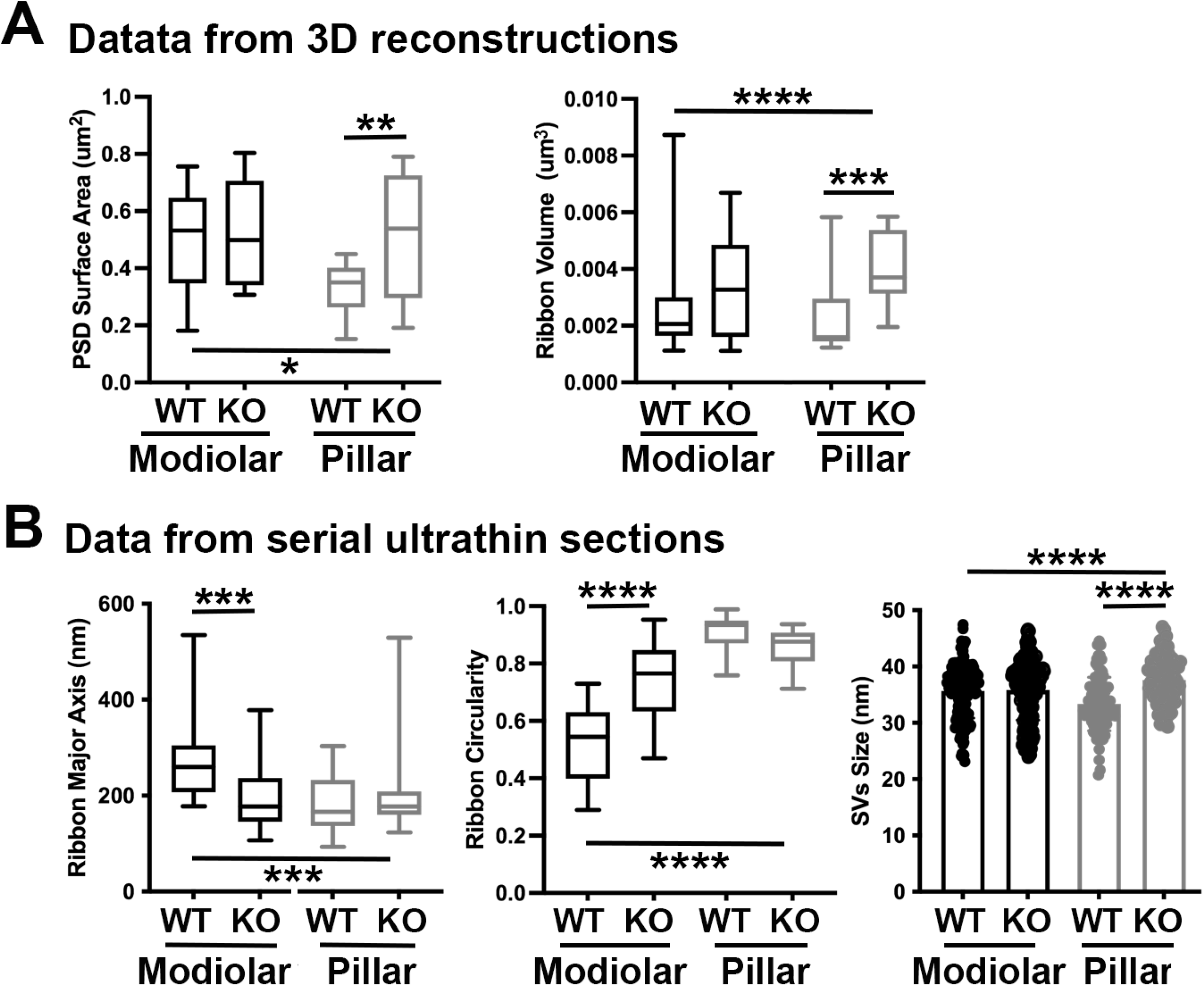
IHC modiolar-pillar structural differences in presynaptic ribbon size, ribbon shape, and vesicle size seen in *GluA3*^WT^ were diminished or reversed in *Gria3*^KO^. **A.** Whisker plots show the quantitative data of the PSD surface area and ribbon volume of *GluA3*^WT^ and *GluA3*^KO^ mice. **B.** Whisker plots of the major axis and circularity of the ribbons of *GluA3*^WT^ and *GluA3*^KO^ mice. Column histogram of the size of SVs of *GluA3*^WT^ and *GluA3*^KO^.

Altogether, our data of 5-week-old male mice show the AMPAR subunit GluA3 is essential to establish and/or maintain the morphological gradients of presynaptic and postsynaptic structures along the modiolar-pillar axis of the IHC. Next, we asked how these early ultrastructural changes in *GluA3*^KO^ correlated with the number of ribbon synapses per IHC and the relative expression of GluA subunits at those synapses.

### An increase in GluA2-lacking synapses precedes a reduction in cochlear output in *GluA3*^KO^ mice

Although *GluA3*^KO^ mice have reduced ABR wave-1 amplitudes at 2-months of age (Garcia-Hernandez et al., 2017), they were not yet different from *GluA3*^KO^ mice at 5-weeks of age (Fig. 1). Given the alterations in ribbon synapse ultrastructure at 5-weeks (Figs. 3-5), we asked if synapse molecular anatomy was also affected in *GluA3*^KO^ mice. Using confocal images of immunolabeled cochlear wholemounts, we analyzed the expression of CtBP2, GluA2, GluA3, and GluA4 at the ribbon synapses between IHCs and SGNs. Visual inspection of the images revealed absence of anti-GluA3 immunoreactivity in *GluA3*^KO^, as well as an obvious reduction in GluA2 labeling and increase in GluA4 labeling relative to *GluA3*^WT^ (Fig. 2; 6A). Despite this, the numbers of paired synapses (CtBP2+GluA2+GluA4) per IHC were similar in the whole cochlea (Fig. 6B; WT: 18.1 ± 2.8; KO: 17.3 ± 3.8; *p* = 0.94, Mann Whitney two-tailed U test) and within apical, middle, and basal cochlear regions (see Fig. 6 caption for more statistical details, Mann Whitney two-tailed U test unless otherwise noted). The numbers of ribbonless synapses per IHC (GluA2+GluA4 – WT: 1.2 ± 0.64; KO: 1.1 ± 0.59; *p* = 0.67) and GluA4-lacking synapses (GluA2+CtBP2 – WT: 0.035 ± 0.044; KO: 0.062 ± 0.095; p = 0.81) were not significantly different. In contrast, the numbers of lone ribbons per IHC (CtBP2-only – WT: 0.97 ± 0.92; KO: 1.6 ± 0.84; *p*= 0.021) and GluA2-lacking synapses (GluA4+CtBP2 – WT: 0.0 ± 0.0; KO: 0.07 ± 0.09; *p*= 0.028) were significantly increased in *GluA3*^KO^ relative to *GluA3*^WT^ (Fig. 6B).

**Figure 6.**
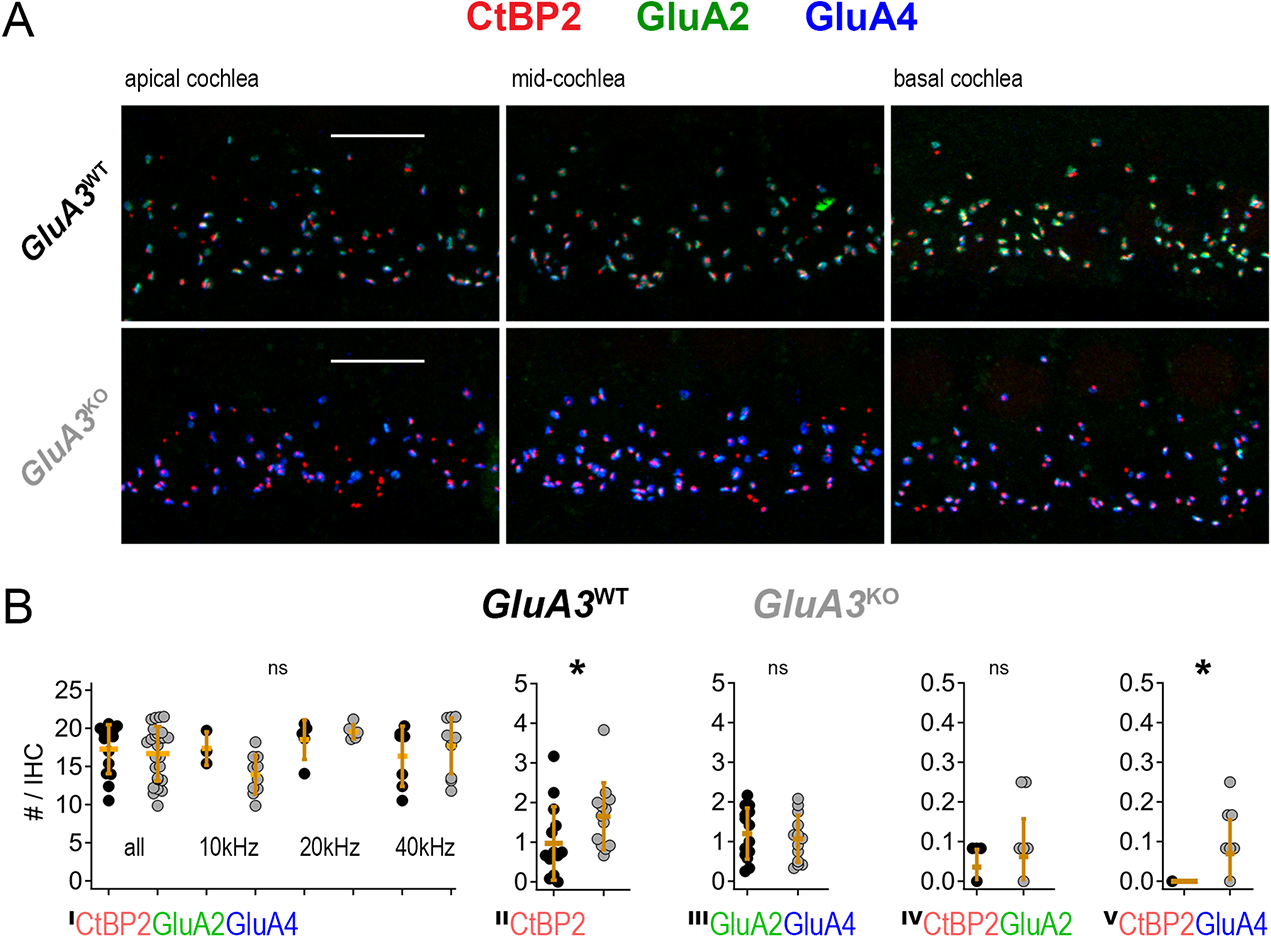
Inner hair cell ribbon synapse counts in 5-week-old male *GluA3*^WT^ and *GluA3*^KO^ mice. **A.** Confocal microscope immuno-fluorescence images of afferent ribbon synapses in organ of Corti whole-mount samples from *GluA3*^WT^ (upper) and *GluA3*^KO^ mice (lower) in the apical, middle, and basal cochlea (left, middle, right). Anti-CtBP2 labels the Ribeye protein in presynaptic ribbons (red); Anti-GluA2 labels the postsynaptic AMPAR subunit encoded by the *Gria2* gene (green); Anti-GluA4 labels the AMPAR subunit encoded by *Gria4* (blue). Each subpanel displays synaptic puncta of approximately 4 inner hair cells. Scale bars: 10 µm. **B.** Quantification of ribbon synapse numbers in images from *GluA3*^WT^ (black: 2,990 synapses; n = 32 images; 5 mice) and *GluA3*^KO^ (gray: n = 2,814 synapses; n = 30 images; 5 mice). Each point represents the mean number of synapses per inner hair cell (IHC) per image; approximately 12 IHCs per image and 6 images per cochlea. **^I^)** Paired synapses per IHC were similar in number for the whole cochlea (*p* = 0.94, U: 484, n_WT_ = 32, n_KO_ = 30) and in each of 3 tonotopic regions centered at 10 kHz (*p* = 0.08, U: 59, n_WT_ = 8, n_KO_ = 10), 20 kHz (*p* = 0.41, U: 61, n_WT_ = 10, n_KO_ = 10), or 40 kHz (*p* = 0.10, U: 42, n_WT_ = 14, n_KO_ = 10; two-tailed Mann-Whitney U test). **^II^)** Lone or ‘orphaned’ ribbons (CtBP2-only) were significantly more frequent in *GluA3*^KO^ (*p* = 0.021, U: 44, n_WT_ = 14, n_KO_ = 13). **^III^)** Ribbonless synapses (GluA2+GluA4) were similar in number (*p* = 0.67, U: 100, n_WT_ = 14, n_KO_ = 13). **^IV^)** Paired synapses lacking GluA4 (CtBP2+GluA2) were similar in number (*p* = 0.81, U: 39, n_WT_ = 7, n_KO_ = 12). **^V^)** Paired synapses lacking GluA2 (CtBP2+GluA4) were significantly more frequently observed in *GluA3*^KO^ (*p* = 0.028, U: 21, n_WT_ = 7, n_KO_ = 12).

### Loss of GluA3 expression reduces synaptic GluA2 and increase synaptic GluA4 subunits

Grayscale and color images (Fig. 2; Fig. 6A) revealed obvious reduction in GluA2 and increase in GluA4 subunit immunofluorescence per synapse, on average, as quantified in Fig. 7A (n = 3 images per genotype). The overall GluA fluorescence per synapse (GluA_Sum_ = GluA2+GluA3+GluA4) tended to be smaller in *GluA3*^KO^ due to the absence of GluA3 subunit fluorescence. Analysis of GluA2 and GluA4 puncta volumes and intensities in one exemplar mid-cochlear image from *GluA3*^WT^ and *GluA3*^KO^ mice revealed that ribbon synapses of *GluA3*^KO^ mice had more compact AMPAR arrays (Fig. 7B) with reduced GluA2 and increased GluA4 fluorescence intensity relative to *GluA3*^WT^ (Fig. 7C). The sublinear Volume vs. Intensity relationship for each GluA subunit suggests synaptic AMPAR density increases with the size of the GluA array in both genotypes (Fig. 7D). Data from three mid-cochlear image stacks from each genotype are summarized in Figure 7E-H (same images as panel A). Relative to the mean of the summed pixel intensities per synapse in *GluA3*^WT^ mice, the overall fluorescence of GluA subunits (GluA_Sum_ = GluA2+GluA3+GluA4) was reduced in *GluA3*^KO^ mice due to the absence of GluA3, despite the much larger increase in GluA4 fluorescence intensity relative to the reduction in GluA2 (Fig. 7E-F). Relative to the mean GluA puncta volume per *GluA3*^WT^ synapse, the mean volumes of GluA2, GluA4, and GluA_Sum_ were all reduced in *GluA3*^KO^ (Fig. 7G). When normalized to the mean puncta volume per image in either group, the distributions of synapse volumes were broadened for GluA2 and GluA4 subunits in *GluA3*^KO^ relative to *GluA3*^WT^ (Fig. 7H). For each image, we calculated the coefficient of variation (CV = SD / mean) in puncta volume for comparison by genotype. The volume of GluA_Sum_ had a CV (mean ± SD, n = 3 images per genotype) of 0.38 ± 0.03 in *GluA3*^WT^ versus 0.51 ± 0.02 in *GluA3*^KO^. For GluA2 volumes, the CVs were 0.39 ± 0.03 in *GluA3*^WT^ versus 0.65 ± 0.065 in *GluA3*^KO^. For GluA4 volumes, the CVs were 0.38 ± 0.04 in *GluA3*^WT^ versus 0.51 ± 0.02 in *GluA3*^KO^. Summed pixel intensity per synapse was more variable than volume and, again, more variable in *GluA3*^KO^ than *GluA3*^WT^. For GluA_Sum_ intensity, the CVs were 0.44 ± 0.04 in *GluA3*^WT^ versus 0.63 ± 0.02 in *GluA3*^KO^. For GluA2 intensity, the CVs were 0.46 ± 0.03 in *GluA3*^WT^ versus 0.76 ± 0.04 in *GluA3*^KO^. For GluA4 intensity, the CVs were 0.45 ± 0.05 in *GluA3*^WT^ versus 0.63 ± 0.02 in *GluA3*^KO^.

**Figure 7.**
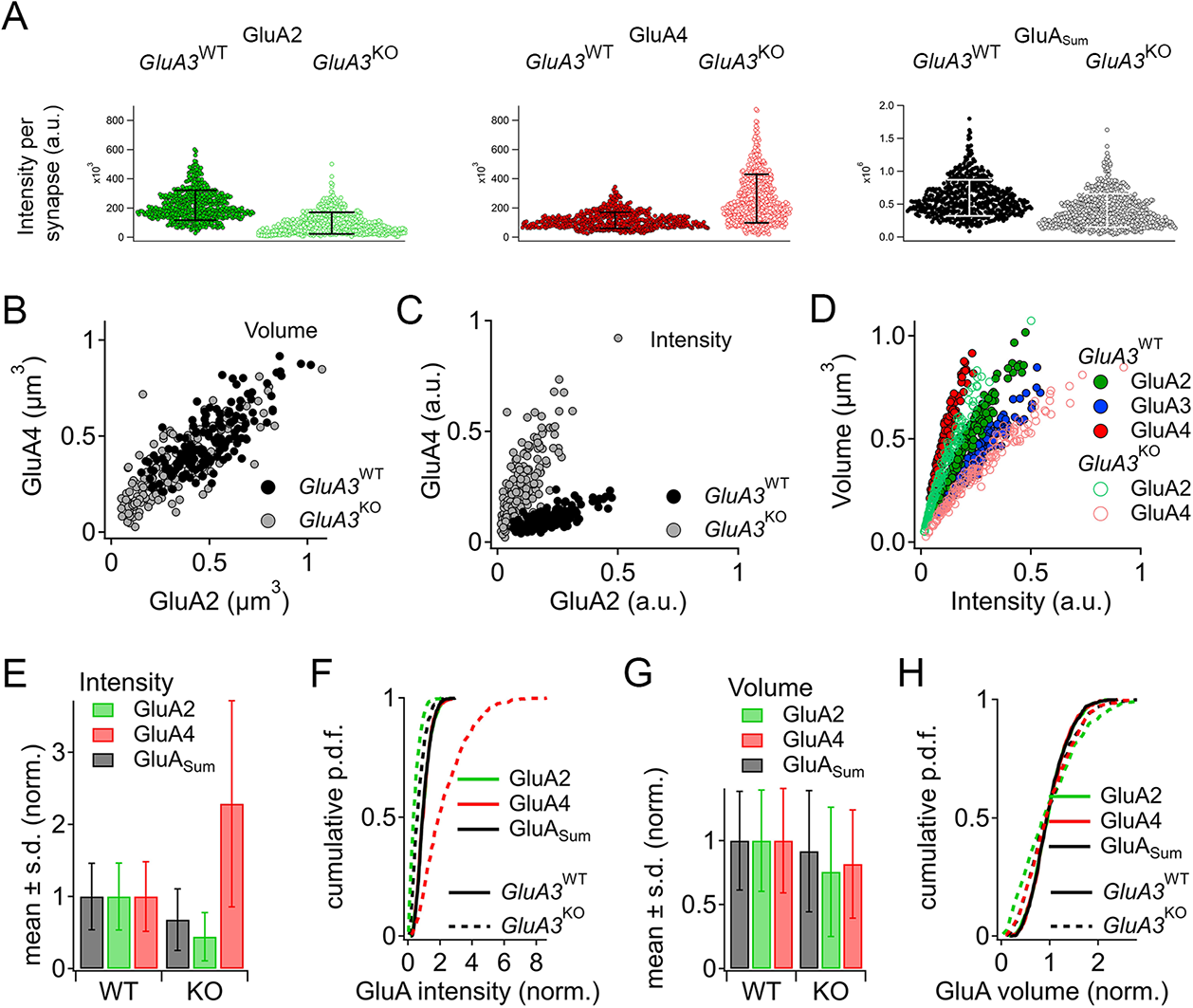
Alteration of AMPAR subunit expression in *GluA3*^KO^ mice. **A.** From images like in Fig. 2: Summed pixel intensity per synapse (raw values, a.u.) for GluA2 (green), GluA4 (blue), and GluA_Sum_ (black). In each subpanel, *GluA3*^WT^ is on left and *GluA3*^KO^ is on right. Bars show mean ± SD; n = 3 mid-cochlear images per genotype, assessed further in panels E-H. **B.** Volume analysis of two exemplar images showing GluA4 vs. GluA2 volume per synapse (µm^3^) in the mid-cochlea of *GluA3*^WT^ (black, n = 148 synapses) and *GluA3*^KO^ (gray, n = 166 synapses). The distribution of GluA4 and GluA2 puncta are shifted to smaller volumes in *GluA3*^KO,^ although the upper ranges are unchanged. **C.** Intensity analysis of the synapses in panel B (summed pixel intensity per synapse) reveals an increase in GluA4 and decrease in GluA2 immuno-fluorescence in *GluA3*^KO^. Intensity values were normalized to the maximum synapse intensity for GluA4. **D.** Volume (µm^3^) vs. summed pixel intensity (norm.) per synapse for *GluA3*^WT^ (filled circles) and *GluA3*^KO^ (open circles) for GluA2, 3, and 4 puncta (green, blue, red). The positive correlation is slightly sub-linear. **E.** Intensity analysis (sum of pixel intensities per synapse) of postsynaptic puncta grouped from *GluA3*^WT^ (n = 545 synapses from 3 images) or *GluA3*^KO^ cochlea (n = 513 synapses from 3 images) shows reduction of overall GluA intensity (GluA_Sum_ = GluA2+GluA3+GluA4, gray) and reduction in GluA2 intensity (green) with increase in GluA4 intensity (red) in *GluA3*^KO^. Data are normalized to the mean WT synapse intensity per group for GluA2, GluA4, or GluA_Sum_. **F.** Normalized data as in panel E displayed as cumulative distributions for *GluA3*^WT^ (solid line) and *GluA3*^KO^ (dashed lines). The overall intensity in *GluA3*^KO^ (black dashed line, GluA_Sum_) is reduced relative to *GluA3*^WT^ (solid black line) due to lack of GluA3 and reduction in GluA2 (green) despite the relatively large increase in GluA4 (red). **G.** GluA puncta volume analysis reveals a reduction of GluA2 and GluA4 volume per synapse in *GluA3*^KO^ relative to *GluA3*^WT^. Data are normalized to the mean WT synapse volume per group for GluA2, GluA4, or GluA_Sum_. **H.** Data in panel G displayed as cumulative distributions. Instead of normalizing to the WT group mean as in panels E-G, here data was normalized to each image mean to visualize differences in the shape of the distributions between *GluA3*^WT^ and *GluA3*^KO^.

To test the statistical significance and to confirm the differences observed in Figure 7 in a larger data set from a replication cohort, we next assessed mean synaptic CtBP2, GluA2, and GluA4 volume and intensity per image in 14 image stacks from each genotype. In image stacks of sufficient quality, we also measured synapse position on the IHC modiolar-pillar axis to sort them into modiolar and pillar groups. Image means and group means are displayed in Figure 8. The volumes of CtBP2, GluA2, and GluA4 puncta were significantly smaller in *GluA3*^KO^ relative to *GluA3*^WT^ (Fig. 8A, all, in µm^3^; CtBP2−*GluA3*^WT^: 0.14 ± 0.02; CtBP2−*GluA3*^KO^: 0.12 ± 0.01, *p* = 0.008 Mann-Whitney U test two-tailed; GluA2−*GluA3*^WT^: 0.47 ± 0.06; GluA2−*GluA3*^KO^: 0.39 ± 0.07, *p* = 0.0001; GluA4−*GluA3*^WT^: 0.45 ± 0.05; GluA4−*GluA3*^KO^: 0.36 ± 0.02, *p* = 4.9e^-6^). In both genotypes, CtBP2 puncta tended to be larger on the modiolar side than the pillar side, on average, but the difference was not significant (*p* = 0.08 for *GluA3*^WT^ and *GluA3*^KO^). In *GluA3*^WT^, GluA2 and GluA4 modiolar-side and pillar-side puncta were not significantly different (*p*= 0.42 GluA2; *p* = 0.23 GluA4). In contrast, in *GluA3*^KO^, GluA2 and GluA4 puncta were significantly larger on the modiolar side than the pillar side (*p* = 0.001 GluA2; *p* = 0.0001 GluA4).

**Figure 8.**
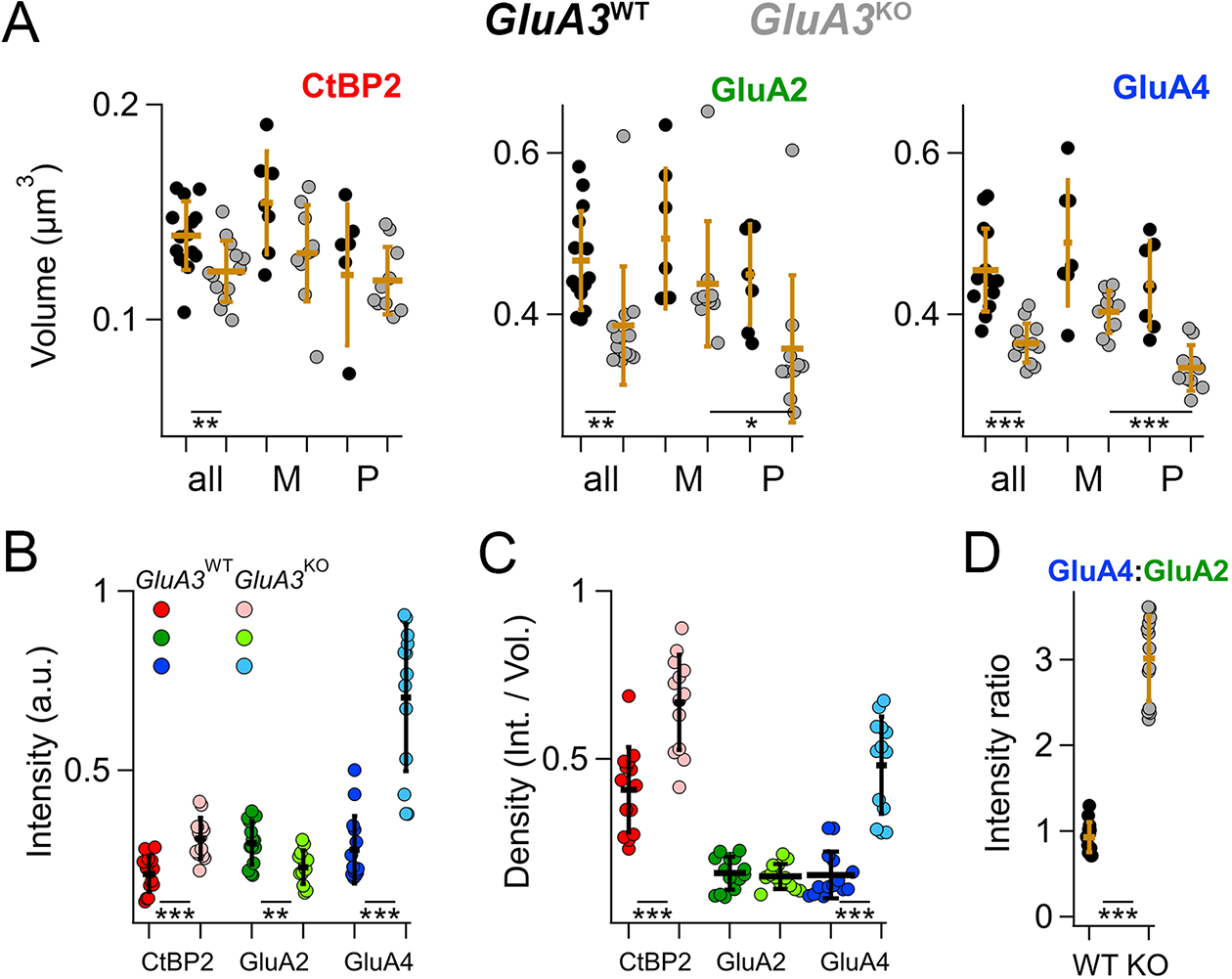
Modiolar-side and Pillar-side Volume, Intensity, and Density of presynaptic ribbon and postsynaptic AMPAR subunits. **A.** Quantification of CtBP2, GluA2, or GluA4 mean volume per image for *GluA3*^WT^ (black, n = 2,990 synapses from 14 images) and *GluA3*^KO^ (gray, n = 2,814 synapses from 14 images). Each point represents the mean number per inner hair cell (IHC) per image; approximately 12 IHCs per image. Gold bars are mean ± SD. For CtBP2, there is an overall reduction in volume in *GluA3*^KO^ (*p* = 0.008, U: 144, n_WT_ = 14, n_KO_ = 13). For GluA2, the overall volume reduction in *GluA3*^KO^ (*p* = 0.0001, U: 168, n_WT_ = 14, n_KO_ = 13) resulted from smaller puncta on the pillar side of *GluA3*^KO^ (*p* = 0.001; U: 90, n_WT_ = 10, n_KO_ = 10). For GluA4, the overall volume reduction in *GluA3*^KO^ (*p* = 4.9e^-6^; U: 176, n_WT_ = 14, n_KO_ = 13) resulted from smaller puncta on the pillar side of *GluA3*^KO^ (*p* = 0.0001; U: 96, n_WT_ = 10, n_KO_ = 10). **B.** Quantification of median intensities per image for data in panel A. CtBP2 intensity increased in *GluA3*^KO^ (*p* = 0.0001; U: 17, n_WT_ = 14, n_KO_ = 13); GluA2 intensity decreased in *GluA3*^KO^ (*p* = 0.01; U: 143, n_WT_ = 14, n_KO_ = 13); and GluA4 intensity decreased in *GluA3*^KO^ (*p* = 5e^-6^; U: 6, n_WT_ = 14, n_KO_ = 13). **C.** Increase in CtBP2 (*p* = 5e^-5^; U: 14, n_WT_ = 14, n_KO_ = 13) and GluA4 median density per synapse (*p* = 5e^-6^; U: 6, n_WT_ = 14, n_KO_ = 13) in *GluA3*^KO^ relative to *GluA3*^WT^. **D.** Increase in GluA4 : GluA2 intensity ratio in *GluA3*^KO^ relative to *GluA3*^WT^ (*p* = 6e^-7^; U: 0, n_WT_ = 14, n_KO_ = 13).

In contrast to mean CtBP2 volume, which was decreased in *GluA3*^KO^ (Fig. 8A), median CtBP2 intensity (Fig. 8B) and density (Fig. 8C) were significantly increased in *GluA3*^KO^. For CtBP2 intensity (a.u.)e^5^ – *GluA3*^WT^: 1.2 ± 0.3; *GluA3*^KO^: 1.8 ± 0.3, p = 0.0001). As shown in the representative images assessed in Figure 7, GluA2 intensity and volume were both reduced (Fig. 8A-B), resulting in no change in GluA2 density (Fig. 8C). For GluA2 intensity (a.u.)e^5^ – *GluA3*^WT^: 1.8 ± 0.4; *GluA3*^KO^: 1.4 ± 0.3, p = 0.01. While GluA4 intensity increased, volume was reduced (Fig. 8A-B), resulting in increased GluA4 density (Fig. 8C). For GluA4 intensity (a.u.)e^5^ – *GluA3*^WT^: 1.7 ± 0.6; *GluA3*^KO^: 4.2 ± 1.2, p = 5e^-6^. Relative to *GluA3*^WT^ synapses, the GluA4:GluA2 intensity ratio was 3x greater on average for *GluA3*^KO^ synapses (Fig. 8D). For GluA4:GluA2 intensity ratio – *GluA3*^WT^: 0.93 ± 0.18; *GluA3*^KO^: 3.0 ± 0.5, p = 6e^-7^.

### Positive correlations between synaptic puncta volumes, intensities, and sphericities in *GluA3*^WT^ are reduced in *GluA3*^KO^ as the range of modiolar-pillar positions is shortened

In *GluA3*^WT^, we commonly observed apparent oscillations in synapse volume as a function of position in the Z-axis of the confocal microscope when the modiolar-pillar dimension was approximately parallel to the Z-axis (Fig. 9A**, lower**). These spatial oscillations were clearer when measured as sphericity (Fig. 9A**, upper**), which was inversely related to volume (Fig. 9A**, right**). We observed a similar phenomenon in *GluA3*^KO^ (Fig. 9B), although the synapses resided in a smaller range along the Z-axis. GluA2 and GluA4 intensities per synapse were positively related in both genotypes (Fig. 9C**, left)**. GluA2 and GluA4 intensities were positively related with CtBP2 intensities, but the relationships were less apparent in *GluA3*^KO^ (Fig. 9C**, center and right**), consistent with the increase in CV measured for GluA2 and GluA4 intensities per synapse in *GluA3*^KO^ relative to *GluA3*^WT^ (Fig. 7). Plotting the GluA4:GluA2 intensity ratio as a function of Z-position revealed that increases of the GluA4:GluA2 intensity ratios in *GluA3*^KO^ tended to be greater for synapses on the pillar side than the modiolar side of the IHC relative to *GluA3*^WT^ (Fig. 9D).

**Figure 9.**
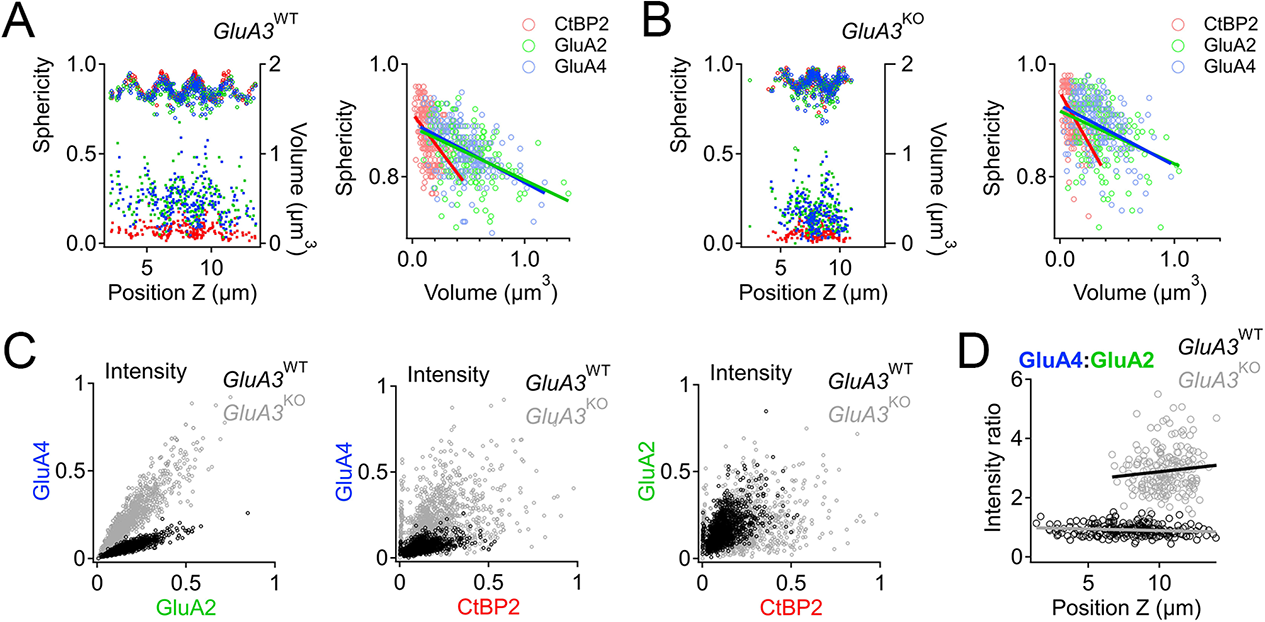
Spatial trends of synapse sphericity, volume, and AMPAR subunit relative abundance in the organ of Corti. **A.** Volume and sphericity per synapse vs. Z-axis-position for an exemplar *GluA3*^WT^ image from Fig. 7 showing spatial oscillations in CtBP2, GluA2, and GluA4. **Right:** Inverse relationship between synapse sphericity and volume for CtBP2, GluA2, and GluA4. **B.** For *GluA3*^KO^, as in panel A. **C. Left:** Normalized intensity of GluA4 vs. normalized intensity of GluA2 for *GluA3*^WT^ (black) and *GluA3*^KO^ (gray). **Center:** Normalized intensity of GluA4 vs. normalized intensity of CtBP2. **Right:** Normalized intensity of GluA2 vs. normalized intensity of CtBP2. **D.** GluA4:GluA2 intensity ratio vs. Z-axis-position. Panels C-D for 6 WT and 6 KO images from the midcochlea.

## Discussion

Hearing depends on the activation of AMPARs on the post-synaptic terminals of auditory nerve fibers (Ruel et al., 1999; Glowatzki and Fuchs, 2002). Cochlear AMPARs are tetrameric heteromers comprised of the pore-forming subunits GluA2, 3, and 4, where the absence of GluA2 results in a CP-AMPAR channel with increased permeability to Ca^2+^ and Na^+^. AMPAR tetramers assemble as dimers of dimers, with the GluA2/3 dimer being energetically favored and prominent (Greger et al., 2019). Our study shows that postsynaptic GluA3 subunits are required for the appropriate assembly of AMPAR GluA2 and GluA4 subunits on mammalian cochlear afferent synapses. Remarkably, we find that GluA3 is also essential for presynaptic ribbon modiolar-pillar morphological distinctions. We propose that postsynaptic GluA3 subunits at IHC-ribbon synapses may perform an organizational function beyond their traditional role as ionotropic glutamate receptors.

### GluA3 is required for appropriate AMPAR assembly at IHC-ribbon synapses

Noise-induced cochlear synaptopathy is caused by excitotoxic overactivation of AMPARs by excessive glutamate release from the sensory inner hair cells (Puel et al., 1998; Kim et al., 2019). Antagonizing the Ca^2+^-permeable subset of AMPARs (CP-AMPARs) pharmacologically can prevent noise-induced synaptopathy while allowing the hearing function to continue through activation of Ca^2+^-impermeable AMPARs (Hu et al., 2020). In the absence of GluA3, GluA2/4 would be the only heterodimer. Homodimers and homomeric tetramers may exist but non-GluA2 subunits preferentially heterodimerize with GluA2 subunits because homodimers are less stable energetically (Rossmann et al., 2011; Zhao et al., 2017). We find that loss of GluA3 alters GluA2 and GluA4 subunit relative abundance, increasing the GluA4:GluA2 ratio (Figs. 8D; 9D), which may increase the number of GluA2-lacking CP-AMPARs at cochlear ribbon synapses of the *GluA3*^KO^ mice. The increase in CP-AMPARs in the *GluA3*^KO^ could make the IHC-ribbon synapses more vulnerable to noise and excitotoxicity as the cochlea matures and ages. In support of this, male *GluA3*^KO^ mice have reduced ABR wave-1 amplitude relative to *GluA3*^WT^ mice by 2-months of age and elevated ABR thresholds by 3-months of age (Garcia-Hernandez et al., 2017). Taken together with previous work, we hypothesize that the 5-week-old *GluA3*^KO^ cochlea may be in a pathological but pre-symptomatic, vulnerable state.

Changes in transcription and mRNA splicing of AMPAR subunits may affect synaptic transmission that can cause excitotoxicity. For example, a decrease in GluA2 and GluA3 *flop* isoforms leads to elevated intracellular Ca^2+^ levels and an increase in the death of retina ganglion cells after glucose deprivation (Park et al., 2016). While there is no evidence at present, the lack of GluA3 could alter the transcription and mRNA splicing of GluA2 and GluA4 in the cochlea during development. In contrast, we find the absence of GluA3 does not alter transcription and mRNA splicing of GluA2 and GluA4 isoforms in the cochleae of 5-week-old *GluA3*^KO^ mice. In both the *GluA3*^WT^ and *GluA3*^KO^ mice, the *flop* splice variant of GluA2 and GluA4 is predominant over the *flip* isoform, as reported in some other CNS glutamatergic neurons (Monyer et al. 1991; Gardner et al., 2001; Pei et al., 2007). Our results argue against the possibility that altered transcription and mRNA splicing of GluA2 and GluA4 lead to excitotoxicity at IHC-ribbon synapses in *GluA3*^KO^ mice (Fig. 1). In addition, we find changes to the presynaptic ribbon suggesting trans-synaptic developmental effects, reminiscent of previous reports on synapse ultrastructure of endbulb synapses in the cochlear nucleus of *GluA3*^KO^ mice (García-Hernández et al., 2017; Antunes et al., 2020). Although young male *GluA3*^KO^ mice have ABR and synapse numbers similar to WT (Figs. 1; 6), we hypothesize these molecular-anatomical alterations to AMPAR subunits result in synapses with increased vulnerability to AMPAR-mediated excitotoxicity that may lead to synapse loss and hearing loss as the mice age in ambient sound conditions.

### Potential trans-synaptic role of GluA3 at IHC-ribbon synapses

Pre- and post-synaptic ultrastructural features of IHC-ribbon synapses are disrupted in the organ of Corti of *GluA3*^KO^ mice (Figs. 3-5). In particular, modiolar-pillar differences are eliminated or reversed in *GluA3*^KO^. Our ultrastructural analysis shows the absence of GluA3 resulted in the loss of the modiolar-pillar difference in PSD surface area seen in *GluA3*^WT^, due to larger PSDs on the pillar side of *GluA3*^KO^ relative to *GluA3*^WT^. Through development, the ribbon-shape changes from largely round to oval, droplet-like, or wedge-like shapes (Wong et al., 2014; Michanski et al., 2019). However, loss of GluA3 resulted in ribbons with shorter long-axes that were more circular due to differences predominantly in the modiolar-side of *GluA3*^KO^ relative to *GluA3*^WT^. This finding is consistent with a developmental defect in a process of ribbon maturation, whereby modiolar-side ribbons become longer and less spherical between 2.5-weeks and 5-weeks of age in C57BL/6 WT mice (Payne et al., 2021). While *GluA3*^KO^ ribbons were shorter in long-axis and more rounded in TEM, they were also more prominent in volume than those from *GluA3*^WT^, suggesting lengthening of the short ribbon axis in *GluA3*^KO^. The increase in ribbon volume measured in *GluA3*^KO^ with TEM was not detected in confocal microscopy, consistent with the ribbon long-axis having the predominant effect on the point spread function when the ribbon short-axis is smaller than the point spread function. Finally, loss of GluA3 eliminated the modiolar-pillar difference in SV size due to an increase in SV size on the pillar side of *GluA3*^KO^. Our ultrastructural data are remarkable and suggest that postsynaptic GluA3 subunits at IHC-ribbon synapses may perform an organizational function beyond their traditional role as ionotropic glutamate receptors. The mechanisms are still unclear, but evidence shows that AMPARs convey a retrograde trans-synaptic signal essential for presynaptic maturation (Tracy et al., 2011). AMPAR subunits may interact with the trans-synaptic adhesion factors Neuroligins and Neurexins (Heine et al., 2008; Hickox et al., 2017). GluA3 is required for the functional development of the presynaptic terminal and the structural maturation of SV size of endbulb auditory nerve synapses in the cochlear nucleus (Antunes et al., 2020). Altered SV size together with a change in the number of AMPARs and their clustering at the synapse contribute to quantal size variation and altered synaptic transmission (Levy et al., 2015). The number of AMPARs at IHC-ribbon synapses is undetermined but with a synaptic surface area ranging 0.1 – 1.5 µm^2^ (Liberman, 1980; Payne et al., 2021), it is estimated several hundred to a few thousand AMPARs at each PSD (Momiyama et al., 2003). In *GluA3*^KO^ mice, we find that despite the decrease in GluA2 and the larger increase of GluA4, the overall intensity and volume for the AMPAR subunits immunolabeling at IHC-ribbon synapses decreases when compared to WT synapses primarily due to loss of GluA3. An increase in relative abundance of CP-AMPARs and a decreased overall abundance of AMPARs in *GluA3*^KO^ are expected to have opposing effects on the size of the synaptic current evoked by glutamate. The ABR wave-1 amplitude is unaltered in male *GluA3*^KO^ at 5-weeks of age suggesting a similar hearing sensitivity to WT mice. However, ABR peak amplitudes are reduced in the male KO at 8 weeks of age (García-Hernández et al., 2017). To strengthen and confirm the potential trans-synaptic role of GluA3 at IHC-ribbon synapses and to compare synaptic strength, further electrophysiological studies need to determine the existence of altered quantal size and quantal content in the *GluA3*^KO^. In the cochlea, afferent synaptic contact formation on the IHC, characterized by a demarcated pre- and postsynaptic density, often precedes ribbon attachment at the presynaptic active zone (AZ) membrane. Ribbon attachment occurs around embryonic day 18 (E18) (Michanski et al., 2019). The presence of postsynaptic AMPARs in those embryonic IHC-ribbon synapses has not been reported. However, patches of GluA2/3 AMPAR subunit immunolabeling were observed during the first postnatal week, juxtaposed to the presynaptic ribbon marker, RIBEYE (Wong et al., 2014). Fusion of ribbon precursors extends after hearing onset and is a critical step in presynaptic AZ formation and maturation (Michanski et al., 2019). This fusion is essential for the functional maturation of afferent synaptic transmission within the cochlea. However, in mice lacking RIBEYE at IHC, it has been shown that features like PSDs, presynaptic densities, and the presence of Ca^2+^ channels and bassoon develop independently of ribbon presence (Jean et al., 2018; Becker et al., 2018). Thus, it is possible that GluA3 plays a direct or indirect role in the recruitment and maintenance of pre- and post-synaptic proteins for example, via its *N*- and *C*-terminus domains. Postsynaptic PDZ domain AMPAR C-terminus interacting proteins such as PSD95 are present at IHC-ribbon synapses early during postnatal development (Tong et al., 2013; Wong et al., 2014). PSD95 interacts with the cell adhesion proteins, neuroligin (Irie et al., 1997; Jeong et al., 2019) which are also detected in the cochlea (Hickox et al., 2017). In the CNS, alignment of postsynaptic AMPARs, PSD95, and Neuroligin-1 together with the pre-synaptic protein RIM (Jung et al., 2015; Krinner et al., 2017; Picher et al., 2017), form a nanocolumn (Tang et al., 2016). The nanocolumns are thought to represent a highly sensitive point in which disruption alters synaptic plasticity, and therefore disrupts synapse function. To understand further the cellular and synaptic mechanisms of hearing and hearing loss, it will be essential to identify which, if any, of the numerous cleft-spanning adhesion systems interact with AMPARs at IHC-synapses, in particular with GluA3.

## Acknowledgments

This work was supported by NIDCD DC013048 (MER) and NIDCD DC14712 (MAR). We thank Nicholas Lozier for his helpful comments on the manuscript.

## Authors’ contributions

M.E.R and M.A.R.: designed research, performed research, analyzed data and wrote and edited the manuscript. M.X. and H.M.C.: performed research and edited the manuscript. A.B. and I.P.: analyzed data and edited the manuscript.

The authors declare no competing financial interests.

